# High-throughput measurement of adipocyte size with open-source software using whole-slide adipose tissue images

**DOI:** 10.1101/2024.10.31.621327

**Authors:** Ramalho Alan, Gauthier Marie-Frédérique, Maltais-Payette Ina, Ostinelli Giada, Hould Frédéric, Biertho Laurent, Tchernof André

## Abstract

The aim of this study was to create and validate a high-throughput method based on open-source software for the measurement of adipocyte diameters in white adipose tissue histological sections. Human omental and subcutaneous adipose tissue samples collected during bariatric surgery were used to prepare hematoxylin and eosin-stained histological slides. Digital images were acquired. Adipocyte diameters were measured both manually and with an automated procedure created using ImageJ. Comparative analysis of our automated method with the manual measurement and associations of the mean adipocyte diameters with cardiometabolic markers were used to validate our method. A total of 377 adipose samples (190 participants) were included in the analysis. Pearson correlation of mean adipocyte diameters shows a strong linear relationship between methods (r=0.88, p<0.0001). The average diameter measured with the automated method was significantly smaller (8.1±5.3µm difference, p<0.0001) compared to the manual method, likely reflecting bias in selecting the cells measured with the manual approach. Pearson correlation analyses between mean omental adipocyte diameters and markers of cardiometabolic risk show that the diameters of both methods are significantly associated with the same parameters (fasting concentrations of TG, HDL-Chol, homeostasis model assessment insulin resistance, and visceral adiposity index values) with no significant differences between methods. There were also no significant differences between the manual and automated method regarding the correlations between mean subcutaneous adipocyte diameters and anthropometric or metabolic markers. In conclusion, we have created and validated a rapid automated method based on open-source software to measure adipocyte diameters from whole-slide adipose tissue images.

## 1 Introduction

The body mass index (BMI) is a poor indicator of individual cardiometabolic risk, and other metrics that better estimate this risk are increasingly favored in clinical practice and in research [1]. In this regard, adipose tissue dysfunction is a major driver of obesity-related cardiometabolic dysregulation [2–5]. Dysfunctional adipose tissue is characterized by reduced adipogenesis, low free fatty acid uptake, impaired triglyceride synthesis, impaired lipid storage, insulin resistance, fibrosis, immune cell infiltration, increased proinflammatory cytokine secretion, and adipocyte hypertrophy [2,4]. It is also strongly associated with visceral obesity, that is, excessive accumulation of fat in the abdominal depots surrounding the internal organs such as the omental (OM) and mesenteric fat depots [2,4], consistent with the notion that visceral obesity is an important risk factor for the development of cardiometabolic dysregulation and cardiometabolic diseases [2,4].

Adipose tissue expands via two main mechanisms: adipocyte hyperplasia where preadipocytes are recruited and develop into small new mature adipocytes through adipogenesis, and adipocyte hypertrophy where existing adipocytes increase in size to accommodate the storage of more lipids [2–5]. Studies have shown that expansion via hyperplasia, typical of subcutaneous (SC) depots, seems to favor the preservation of cardiometabolic health [2,3]. Conversely, expansion via hypertrophy is an important indicator of adipose tissue dysfunction and increased cardiometabolic risk [2,3]. Indeed, adipocyte hypertrophy is often implicated in visceral obesity and has been shown to be associated with an increased secretion of proinflammatory factors as well as with altered lipid metabolism, impaired glucose-insulin homeostasis, ectopic fat accumulation, and cardiovascular disorders, independently of BMI [2–4,6].

Consequently, measuring adipocyte size can provide indirect information on adipose tissue function, fatty acid and glucose metabolism, and cardiometabolic pathophysiology [4,6,7]. Analysis of adipocyte size can also help better understand the pathophysiological processes of obesity and cardiometabolic diseases as well as to characterize treatment responses to dietary interventions, nutraceuticals, bioactive compounds, and medication [8,9].

Currently, there is no gold standard method to measure adipocyte size [4,6]. There are three main methods utilized in research: imaging of collagenase digestion-isolated adipocytes, sizing of osmium tetroxide-fixed cells, and histomorphometry [3,4]. Histomorphometry involves analyzing hematoxylin and eosin (H&E)-stained histological slides of adipose tissue using an optical microscope to measure the diameter or cross-sectional area of the adipocytes [4]. This method is considered to be the most reliable, but it still has drawbacks [8]. First, there are several inherent assumptions including a uniform cell size distribution and that cells are showing their largest diameter or cross-sectional area [4]. Furthermore, fixation of the tissue reduces volume of the cells by approximately 15-20%, directly measuring the diameter is not reliable for irregularly shaped cells, distortion from poor fixation can occur, crush artifacts can be introduced during processing, and the quality of the stain can vary between laboratories [4,6,8]. Differences can also exist regarding specificities in cell sizing protocols which may influence the final results and the ability to compare results between laboratories [10,11]. The most important drawback, however, is that this method is very labor-intensive, time consuming, and has a high risk of measurement and observer bias [7,8,12]. Due to the time required, only 100 to 300 cells per sample are typically measured, decreasing measurement accuracy [12,13]. These are inherent issues of manual histomorphometry as it requires the measurer to visually identify each adipocyte in the tissue section and then measure their diameter or cross-sectional surface area. With the advent of biological image analysis software, such as ImageJ (National Institutes of Health, USA), the measurement of adipocyte surface area has become easier. ImageJ is a free easily accessible open-source scientific image processing and analysis software platform originally created by the National Institute of Health of the United States [14]. ImageJ is relatively easy to use and has been widely adopted by the scientific community.

To address the main drawbacks of the histomorphometric method, several teams have developed automated programs or protocols to detect and measure adipocyte size using histological slides [7–9,11,12,15–18]. However, many of these automated solutions have issues such as adaptability, cost, reliability, accessibility (both short- and long-term), and usability. Many tools cannot be adapted sufficiently to account for considerable procedural differences such as sample quality preservation, paraffin inclusion, microtomy, staining, and even imaging [10]. The cost of paid software solutions as well at the costs of the equipment required can be prohibitory [12]. Some software tools have been discontinued or are not accessible through secure channels [7,9,17,18].

Many of the current software solutions typically analyze digital photomicrographs of the tissue sample, limiting the number of adipocytes measured [9,12,13]. Photomicrographs are used for manual analysis as manually measuring each adipocyte in a complete section would require an exorbitant amount of time. Cell sizing software tools should be capable of analyzing whole-slide images of the tissue samples to maximize the number of adipocytes measured [12]. However, many of the free software options were not designed to do this due to file size restrictions. Recent advancements in both file size capacity of software tools and the affordability of slide scanners for the acquisition of high-quality whole-slide images have opened the door for significant progress in this regard [12].

Ultimately, there are important issues with the current histomorphometric methods of fat cell sizing. The aim of this study was to develop and validate an easy-to-use method to measure adipocyte size in whole tissue images. We hypothesized that measurements can be made automatically and that average adipocyte diameter measurements obtained with the automated method are consistent with those obtained with the manual method, while requiring significantly less time and measuring a larger number of cells per sample. Additionally, we hypothesized that the results obtained with the automated method are equivalent or superior to those of the manual method for the prediction of cardiometabolic risk markers.

## 2 Results

### 2.1 Manual and automated cell size measurements

Regarding the initial elaboration phase of our study, the average adipocyte diameters from 16 samples obtained with the manual and automated methods were compared for the purpose of establishing measurement parameters. Spearman correlation analysis showed a strong significant association between the methods (r=0.90, p <0.0001). Paired t-test analysis indicated that the average adipocyte diameters obtained with the two methods were not significantly different (p=0.10) (data not shown).

For the validation study, a total of 370 and 374 samples (for the manual method and automated method, respectively) from 190 participants were included in the analysis (**Table 1 and Table 2**). As shown in **Table 2**, the average adipocyte diameter was 77.1 ± 11.0 µm and 69.1 ± 10.5 µm for the manual and automated method, respectively. These average diameters were shown to be significantly different by paired t-test (p<0.0001) (data not shown). The approximate amount of time dedicated to the manual analysis was 10 minutes per tissue compared to 0.8 minute per tissue for the automated method (based on a subgroup of 300 samples). The automated method detected an average of 878 ± 511 cells compared to 130 ± 34 cells for the manual analysis (p<0.0001).

**Table 1:**
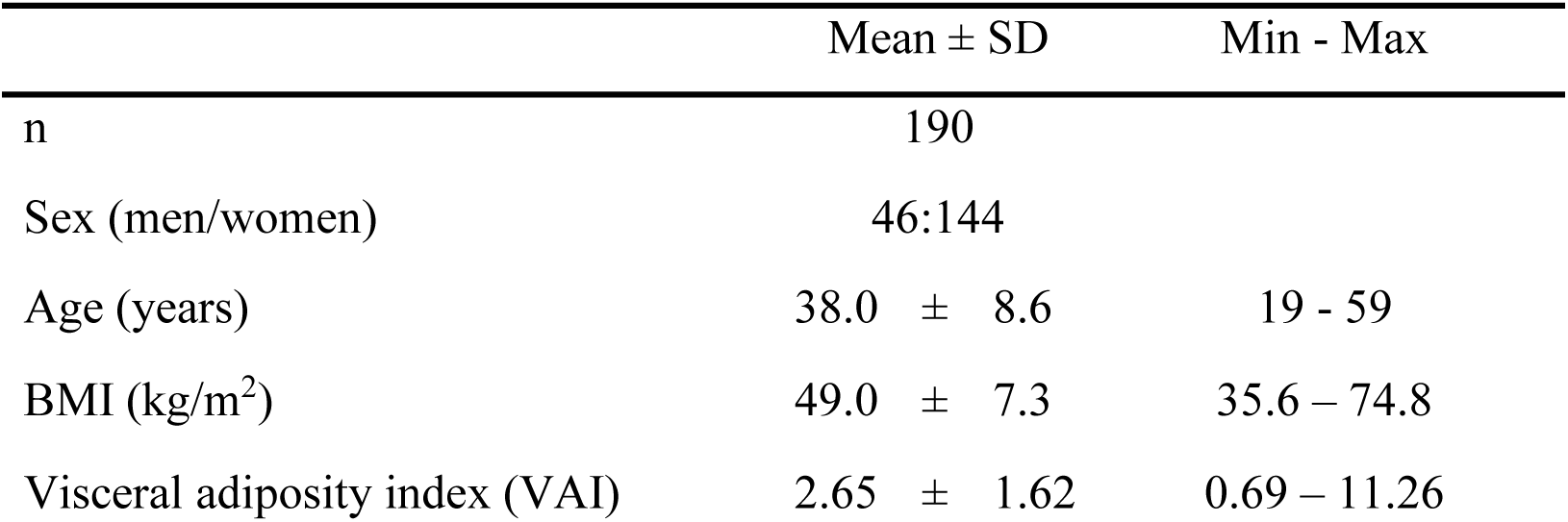
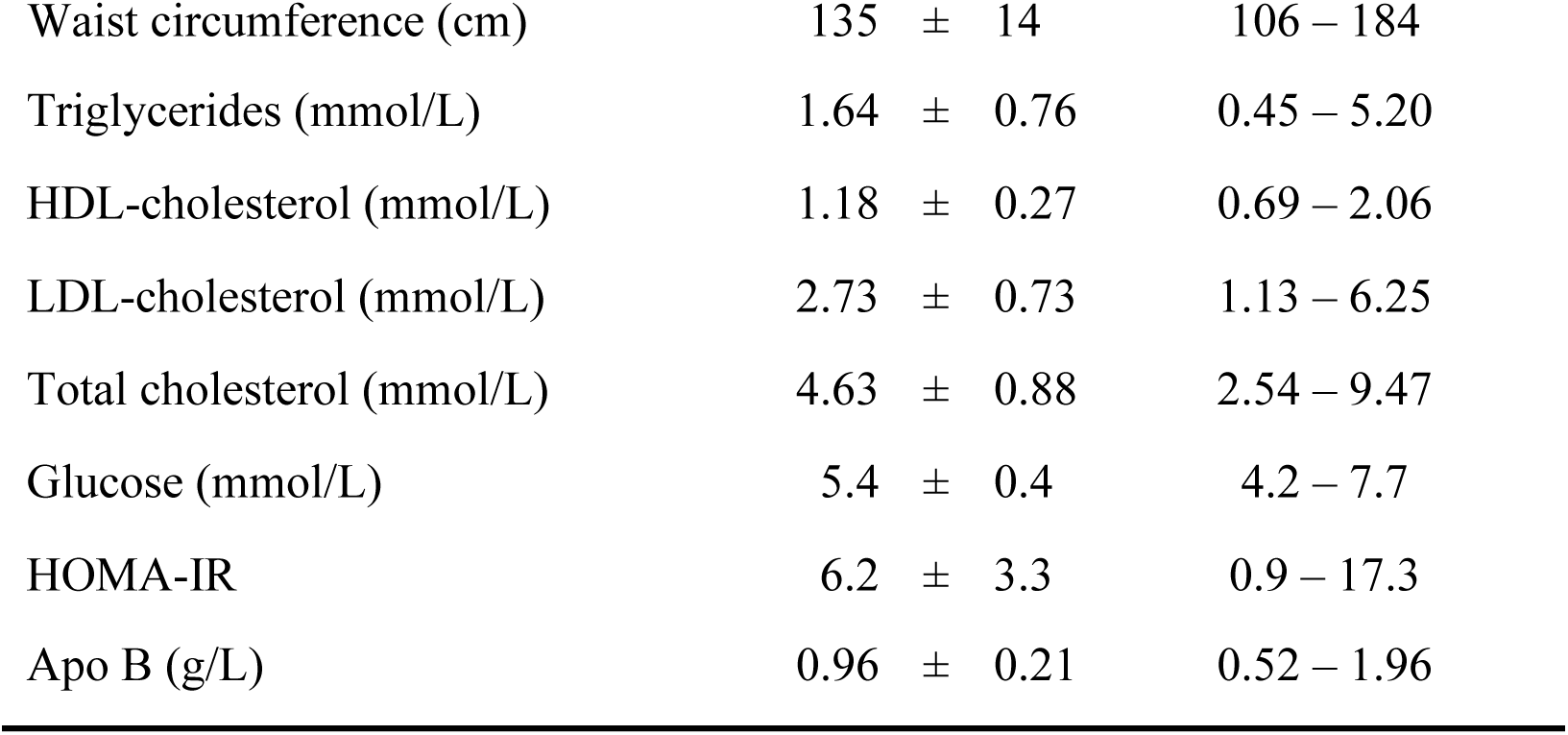
Participants’ anthropometric characteristics and metabolic markers levels. Data are expressed as the mean ± SD and minimum and maximum values.

**Table 2:** Summary results of all adipocytes diameter (OM and SC) with the manual and automated method. Data are expressed as the mean ± SD for the measured cell number and cell diameter. Time required for analysis includes file processing and handling.

|  | Manual method | Automated method |
| --- | --- | --- |
| N | 370 | 374 |
| Time required for analysis | ~ 10 minutes/section | ~ 0.8 minutes/section |
| Average number of cells measured ±SD | 130 ± 34 cells | 878 ± 511 cells |
| Average cell diameter ± SD | 77.1 ± 11.0 $\mu\text{m}$ | 69.1 ± 10.5 $\mu\text{m}$ |

### 2.2 Comparisons between manual and automated cell size measurements

A Pearson correlation analysis was performed between the manual and automated cell size (for the two depots) and is presented in **Figure 1a**. The Pearson coefficient of r=0.88 indicates that the two methods are strongly consistent. We observed in the Bland Altman plot in **Figure 1b** an average difference of −8.10 µm between the mean automated and manual cell diameters, as well as limits of agreement of 2.22 µm and −18.43 µm. Most of the data points are within the limits of agreement, indicating good agreement between methods. The magnitude of the difference does not appear to vary based on the average diameter, suggesting the absence of systematic bias related to the magnitude of the measurement [19]. **Figures 1c and 1d** show that the average cell size measured with the automated method is significantly smaller compared to the manual method in both the visceral and SC depots (−9.55 µm, p<0.0001 and −6.61 µm, p<0.0001, respectively) and this is also true for the majority of individual samples.

**Figure 1:**
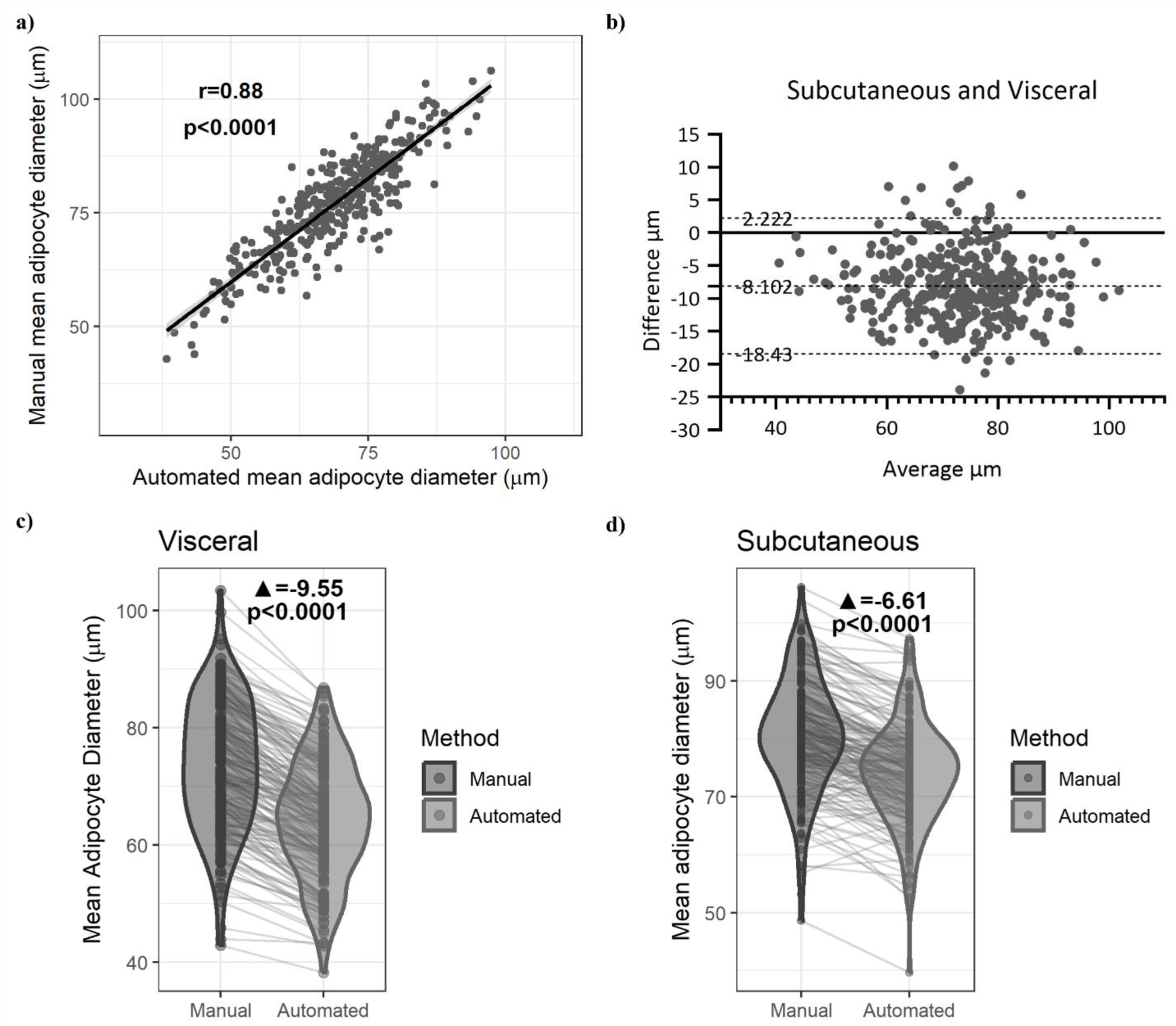
Comparison of adipocyte diameters measured by the two methods. Pearson correlation in all cell diameter (OM and SC) between the manual and automated method **(a)** and Bland-Atlman plot with size differences **(b)**. Paired t-test between the two methods in omental **(c)** and subcutaneous **(d)** adipose tissues.

### 2.3 Associations with metabolic markers

The participants for this study were on average 38.0 ± 8.6 years old, including 46 men and 144 women with an average BMI of 49.0 ± 7.4 kg/m^2^ and an average waist circumference of 135 ± 14 cm (**Table 1**). Pearson correlation analyses were performed to evaluate the association between cardiometabolic health markers and the mean adipocyte diameter obtained with each measurement method (i.e. manual and automated) for each adipose tissue depot. The results are presented in Figure 2. The automated cell size method was not significantly different from the manual method for any of the cardiometabolic health markers for either depot, as determined using Meng’s Z-test (data not shown).

**Figure 2:**
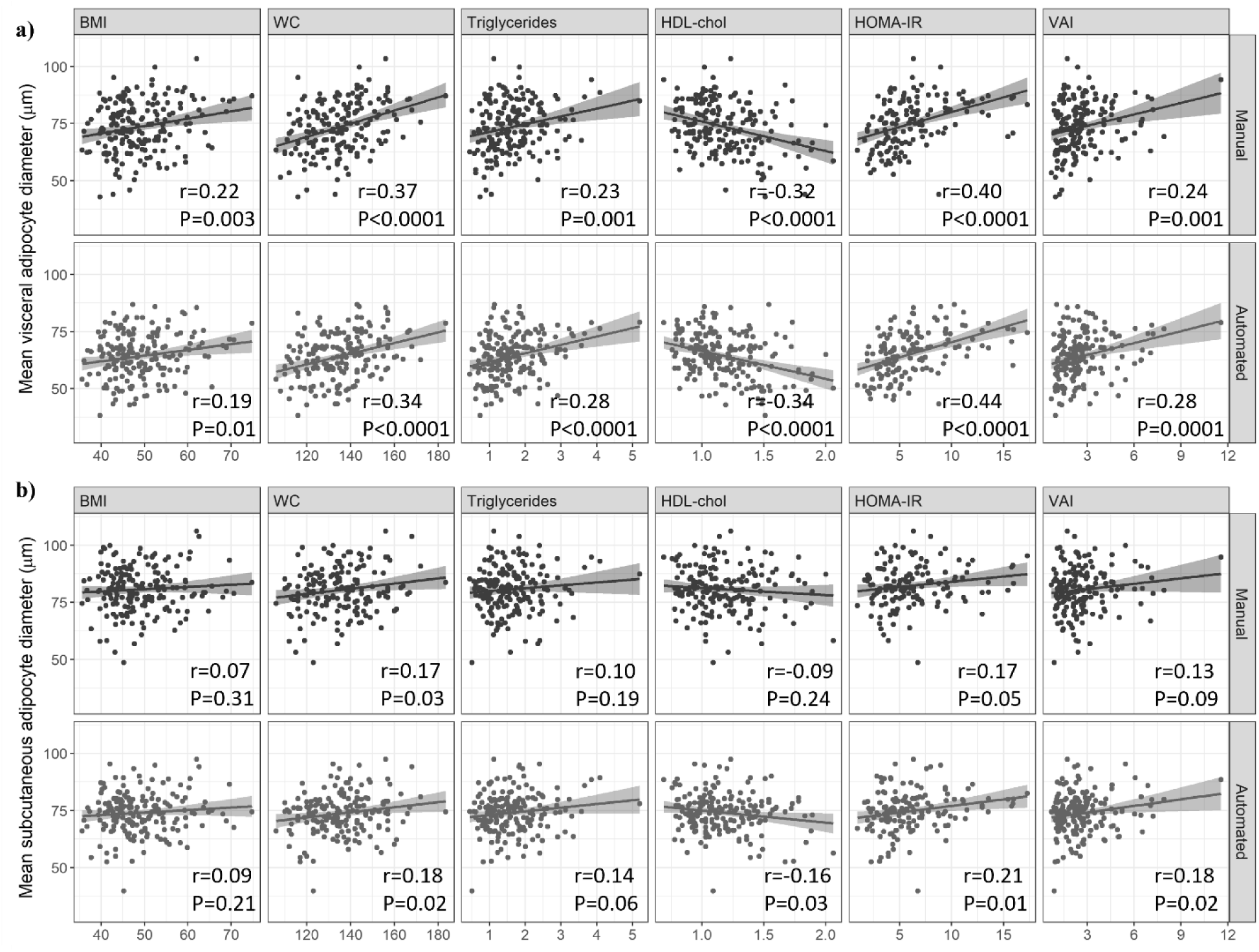
Comparison in association with anthropometrics and metabolic markers. Pearson correlation between mean adipocyte diameter with BMI, WC and cardiometabolic markers (TG, HDL-Chol, HOMA-IR and VAI) between the two methods in omental **(a)** and subcutaneous depot **(b)**.

Regarding the OM depot, we observed in Figure 2a, although not significantly different, a slightly stronger correlation and lower p-value with the automated method with triglyceride (TG) levels (r=0.28, p<0.0001 vs r=0.23, p=0.003), HDL-Cholesterol (HDL-Chol) (r=-0.34, p<0.0001 vs r= −0.32, p<0.0001), the homeostatic model assessment of insulin resistance (HOMA-IR) (r=0.44, p<0.0001 vs r=0.40, p<0.0001), and the visceral adiposity index (VAI) (r=0.28, p=0.0001 vs r=0.24, p=0.001) compared to the manual method.

Regarding the SC depot (Figure 2b), although not significantly different, a slightly higher correlation strength was observed with the automated method for the association with HDL-Chol (r=-0.16, p=0.03 vs r=-0.09, p=0.24), HOMA-IR (r=0.21, p=0.01 vs r=0.17, p=0.05) and VAI (r=0.18, p=0.02 vs r=0.13, p=0.09) compared to the manual method. Notably, the association with HDL-Chol and VAI became significant with the automated method. No significant associations were observed with total Cholesterol (Chol), LDL-Chol, and apolipoprotein B (apoB) with either method in either depot.

## 3 Discussion

Our analysis shows that our automated method for measuring adipocyte size is rapid, valid, and reliable. Pearson correlation analysis shows a strong agreement between both methods for the determination of mean adipocyte diameters. Pearson correlation analyses and Meng’s Z-test also demonstrate that the associations between different cardiometabolic health markers and mean adipocyte diameters measured using the automated method were equivalent to the associations with the manual method. Adipocyte size is often used in research as a marker of adipose tissue function and cardiometabolic health. As such, the significant associations between other cardiometabolic health markers and the automated mean adipocyte diameter and their agreement with the associations with the manual mean adipocyte diameters clearly demonstrate the validity of our method. It should be noted that the weight status of the participants in this study likely influenced these associations. Expansion through hypertrophy has its limitations as adipocytes can only increase in volume up to a certain point [2,4]. Mean fat cell size eventually plateaus in individuals with large adiposity excess as can be the case with candidates of bariatric surgery [2,4,11]. This plateauing weakens the ability to detect significant associations with other variables [4]. This may explain the lack of significant associations with some of the cardiometabolic risk markers and why the associations observed with BMI and WC were weaker than expected based on data from other studies [3,13].

Despite the strong association between methods, the paired t-test, grouped violin plots, and the Bland-Altman plot demonstrate that the automated mean adipocyte diameters are on average significantly smaller than those of the manual method. The average difference between the two methods is 8.1 µm. Because there is no gold standard method for cell size measurement, it is possible that this difference reflects better estimation of average cell size by our automated method compared to the manual method. The manual method is dependent on the judgement of the user in both the selection of the tissue section subregions to use and the identification of adipocytes to measure. The selection of subregions is ideally completely randomized, but many factors such as section edges, damaged subregions, and fibrotic subregions can influence this selection. This may result in preferentially identifying better-looking subsections. Large adipocytes may be easier to notice and identify visually than smaller ones, which could result in an overestimation of average size with the manual approach. The automated method targets the entire section, thereby eliminating such biases. Analyzing the entire histological section instead of a few subregions also greatly increases the number of cells measured for each sample. Increasing the number of cells measured increases the likelihood that the estimation of average cell size is accurate [12]. Other teams have obtained similar differences in mean adipocyte size. Maguire et al (2020) and Palomäki et al (2022) [11,12], both observed that mean cell size was smaller when using an automated method with whole-slide images compared to methods that used photomicrographs of tissue sections. Interestingly, Maguire et al (2020) [12] only observed this in high-fat diet mice, and the opposite was true for chow-fed mice. Like us, Palomäki et al (2022) [11] used adipose tissue samples from patients with obesity, but the difference they observed between sizing methods was much smaller at −0.8 µm. This may be due to methodological differences between their study and ours as they measured approximately 300 cells per sample using photomicrographs while also using a smaller number of tissue samples [11]. Due to these reasons and the significant correlations with cardiometabolic risk markers previously discussed, it is possible that our automated method produces a more reliable estimation of mean adipocyte size compared to the manual method.

Our method has several advantages compared to the manual method. As discussed, our automated method is significantly faster and uses whole-slide images of H&E-stained tissue sections, significantly increasing the number of adipocytes measured. The automated detection of adipocytes reduces the potential of observer and measurement bias. Our method is also very easy to implement as its main component is a macro for ImageJ, that we provide here and can easily be copied, making it very accessible. The only software needed is ImageJ, which is free, easily accessible, and can be run on low-powered computers. The fact that ImageJ was developed by the NIH and that it has a dedicated userbase provides good assurance that it should be maintained for the foreseeable future. ImageJ is also easy to learn, especially for this application.

As mentioned, other teams have developed automated protocols using software, but many of these solutions have issues including software cost, short- and long-term accessibility, and the inability to use whole-slide images. Notable exceptions to most of these issues are Adiposoft, a plugin created for ImageJ, and QuPath, another free open-source software for the analysis of digital biological images [8,12,20]. Adiposoft was created and validated to analyze the number and size of white adipocytes from adipose tissue histological sections [8,21]. Originally, it only analyzed photomicrographs but is now capable of analyzing whole-slide images. Indeed, Adiposoft is a capable program solution, but we encountered issues when attempting to use it with our samples. While Adiposoft analyzed many of our samples without issue, there were significant problems with some samples such as adipocytes not being detected and the identification of large empty areas as adipocytes. Our experience suggests that it may be difficult to adapt Adiposoft, using its current settings options, to certain experimental conditions. Adiposoft also outputs a unique spreadsheet for each image, complicating data management when analyzing many samples. QuPath is a powerful tool for the analysis of whole-slide images and large 2D data [20]. Maguire et al (2020) developed an interactive plugin for QuPath for the measurement of adipocytes [12]. Their results show that their method reliably measures adipocyte number and size from whole-slide images [12]. However, their method involves several steps that require both QuPath and ImageJ. The adaptability of their method to different experimental conditions is also uncertain. The QuPath method developed by Palomäki et al (2022) only uses tools that are already built into QuPath and no other analysis software is required. Their analysis shows that their method is valid for good and average quality whole-slide images and photomicrographs [11]. However, the method requires training a pixel classifier with good quality histology sections of adipose tissue [11]. This step increases the complexity and requires technical knowledge, which may hinder other teams from adopting the method. Ultimately, even with free software solutions capable of analyzing whole-slide images, there are issues that hinder their adoption. In contrast, our method is more straightforward and simpler to implement. It requires only ImageJ and our macro, which is arguably easier. Our macro uses standard image processing methods, which should allow other teams to easily understand it and adapt it to their experimental conditions. Furthermore, the macro does not need to be downloaded, maximizing its accessibility.

Our study has several strengths. First, to validate our automated method, we used both correlation analyses and Bland-Altman plots to evaluate the strength of the linear relationship and agreement between the methods, respectively. While a strong correlation between methods of measurement of the same variable is expected, it does not indicate that there is good agreement between methods [19]. Bland-Altman plots evaluate the differences between the methods, allowing for a better determination of comparability between methods [19]. Second, 367 white adipose tissue samples obtained from male and female human participants were analyzed by both methods. Such a large sample size greatly increases the validity of our analysis and is much larger than samples used by other teams in the validation of their cell-sizing tools [12,15,16]. Furthermore, we used both SC and OM adipose tissue samples from patients from two different cohorts, which indicates how our automated method can be applied to different scenarios. Third, we evaluated how the methods influence the associations between cardiometabolic health markers and mean cell diameters to establish the validity and usefulness of our automated method.

This study does have some limitations. First, all the samples came from participants with obesity undergoing bariatric surgery. As such, it cannot be stated for certain that our method would adequately perform in populations without obesity. This is unlikely to be an issue due to obesity being a very heterogeneous disease and that we used a very large sample size which included varied types of cellular morphologies. Rodent white adipose tissue is similar to that of humans regarding morphology and gross anatomical regions, indicating rodent tissues could likely be analyzed with our method [17]. Second, all the histological slides were prepared by the histology department at our Institute by highly trained technicians using high-quality equipment. The slides were typically of high quality. As such, our method may not function well with lower quality slides. However, poor quality slides are an impediment to all sizing methods, and, as mentioned, our macro can be adapted to optimize its performance to different experimental conditions. Our method also requires the use of whole-slide images with uniform illumination, but the recent decrease in the cost of slide scanners alleviates this issue [12]. Third, only mean adipocyte diameter was considered and not adipocyte size distribution in each sample. As only approximately 100 adipocytes were measured per sample, it is possible that some size subgroups would be underrepresented with the manual method, resulting in the comparison of the two methods in this regard to be of questionable relevance.

In conclusion, we have successfully developed a rapid, automated method for the measurement of adipocytes in H&E-stained histological sections of white adipose tissue. This automated method consists of a straightforward macro for ImageJ that analyzes whole-slide images of histological white adipose tissue sections. We have shown that this method is valid and reliable. Compared to the usual manual method, our automated method allows for the measurement of a much greater number of cells while requiring considerably less time. Furthermore, our method can easily be adopted and adapted by other teams, which could increase the uniformity between laboratories.

## 4 Methods

### 4.1 Participants characteristics and tissue sampling

Participants are patients who underwent bariatric surgery and were recruited through the Biobank infrastructure at the *Insititut universitaire de cardiologie et de pneumologie de Québec – Université Laval*. All participants provided written, informed consent. This study was approved by the medical ethics committees of Laval University (approval numbers 2017-2710, 2021-3398 and 2019-3218) and adheres to the principles of the Declaration of Helsinki. Clinical and anthropometric characteristics such as age, sex, height, weight, BMI, and waist circumference were recorded during the pre-surgery consultation or on the morning of surgery. Fasting blood samples were obtained and lipid profiles (total Chol, HDL-Chol, LDL-Chol, TG, and apoB levels) and hematological parameters (fasting insulin, fasting glucose, and HbA1c) were measured by the biochemistry department of the Institute. OM and SC adipose tissue samples were collected from the greater omentum and the abdominal SC adipose tissue compartment, respectively, during surgery and immediately frozen at −80°C. Approximately 60 to 80mg of frozen adipose tissue was fixed in 10% formalin (Sigma-Aldrich®, HT501128-4L) overnight at 4°C and embedded in paraffin the following day in accordance with our validated protocol [10]. Five µm sections were mounted on slides, stained with H&E and used for the manual and automated measurements of cell size. HOMA-IR was calculated according to the equation: fasting insulin (µU/mL) * fasting glucose (mmol/L) /22.5 [22]. VAI was calculated using validated sex-specific equations [23]. The equations are as follows: males: 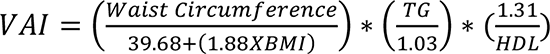 females: 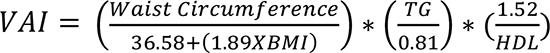[23]

Initially, a small group of 16 samples were selected for the elaboration of the automated method and an initial validation. The samples were obtained from four participants (two men and two women, 45 ± 9.7 years old, BMI 48 ± 6.6 kg/m^2^ at the time of bariatric surgery). This group allowed us to determine the parameters for the automated measurements in a variety of cell size ranges using adipose tissue samples from two depots.

Subsequently, validation of the method was performed with a larger group composed of 190 participants who had undergone bariatric surgery and were included in this study.

### 4.2 Manual adipose cell size measurements

Briefly, five subregions with minimal damage and artifacts were randomly acquired from each histological adipose tissue section using a Leica DMLB microscope (Leica, Wetzlar, Germany) coupled with a MotiCam580 (Motic, British Columbia, Canada) at 5x magnification for each sample. Manual histomorphometric analyses were performed blinded to patient identity or type of depot according to the method described by Laforest et al (2018) [10]. The digital photomicrographs were converted to 8-bit, the background was subtracted, background noise was reduced using a median filter, and a manual threshold was applied to convert the images to binary in order to segment adipocytes using ImageJ (National Institutes of Health, USA). The cross-sectional area of each adipocyte was manually measured using the wand tool on each image. Incomplete cells on the edge of the images were excluded. The paintbrush tool was used to smooth the cell surface or fill-in membranes when needed. A minimum of 100 cells per tissue were measured. The cross-sectional areas were used to calculate diameters, expressed in µm.

### 4.3 Automated adipose cell size measurements

Whole-slide images were acquired with an Axio Scan Z1 slide scanner (Zeiss, Oberkochen, Germany) at 20x magnification. Whole-slide digital images were first converted to 8-bit images and then into binary using ImageJ. The thresholding method was selected based on the best segmentation of the adipocytes. Multiple thresholding methods were tested and triangle thresholding produced the best result for our images. The automatic thresholding compensates for differences in light intensity during acquisition of H&E staining variation within samples. The threshold is adjusted and optimized automatically on each image. Other parameters such as removing outliers, eroding, and despeckling were tested to reduce background noise.

Area analysis was performed through a particle analysis component in ImageJ. The particle analysis is based on the size and the circularity of objects. Minimum and maximum size cut-offs of 315 and 40000 µm^2^ (equivalent to a diameter of 20 and 225 µm), respectively, were applied so all objects inferior to 20 µm and superior to 225 µm in diameter were excluded. The minimum cut-off enabled the exclusion of objects unlikely to be adipocytes. The maximum cut-off is important to control for potentially damaged tissues. Objects larger than 225 µm in diameter likely represent two or more cells where the cell membrane collapsed or was damaged during tissue processing. This would result in an overestimation of cell size. The roundness of mature adipocytes allows us to use circularity as an important component of this analysis. Different circularity value cut-offs were tested during the initial elaboration and validation phase. The circularity cut-off values have been selected to maximize the number of detected cells while having the best fit with the manual cell sizes without including false detections due to poor tissue quality (cell damage). Once the steps of the procedure were established, the different parameters were incorporated into a macro file in ImageJ to perform automatic fast-paced analysis on all images. The macro is as follows:

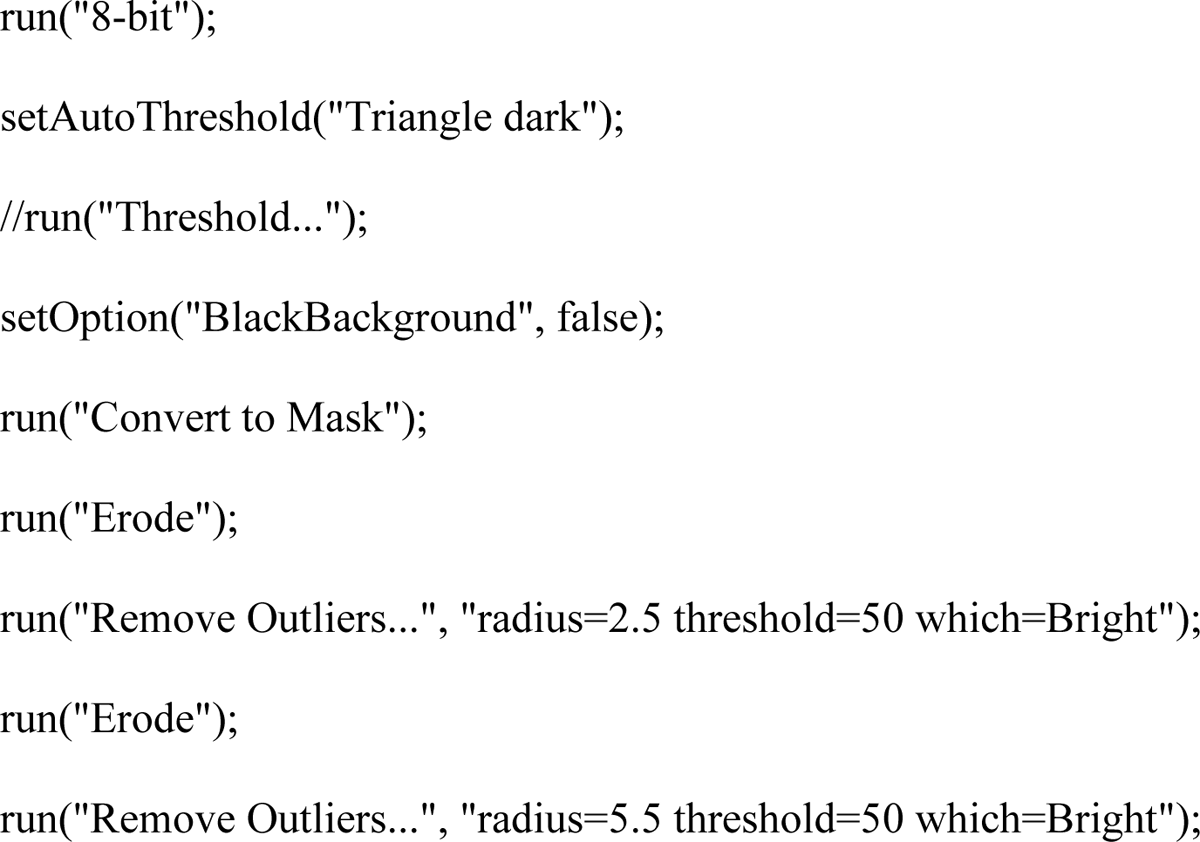
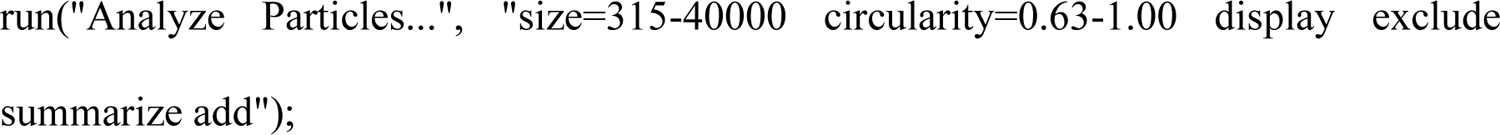

Mean surface areas were extracted and converted to diameters expressed in µm. For quality assurance, when the number of detected cells was less than 200, the whole-slide image was visually inspected to determine why the count was low. If there was an issue in cell detection, the threshold was manually adjusted to improve segmentation and the image was reanalyzed. If the detection of cells appeared accurate, the initial results were kept if the total number of cells detected was greater than 100. If less than 100 cells were detected, this was considered insufficient, and the results were omitted from subsequent analyses. A simplified schematic overview of the two methods is presented in Figure 3.

**Figure 3:**
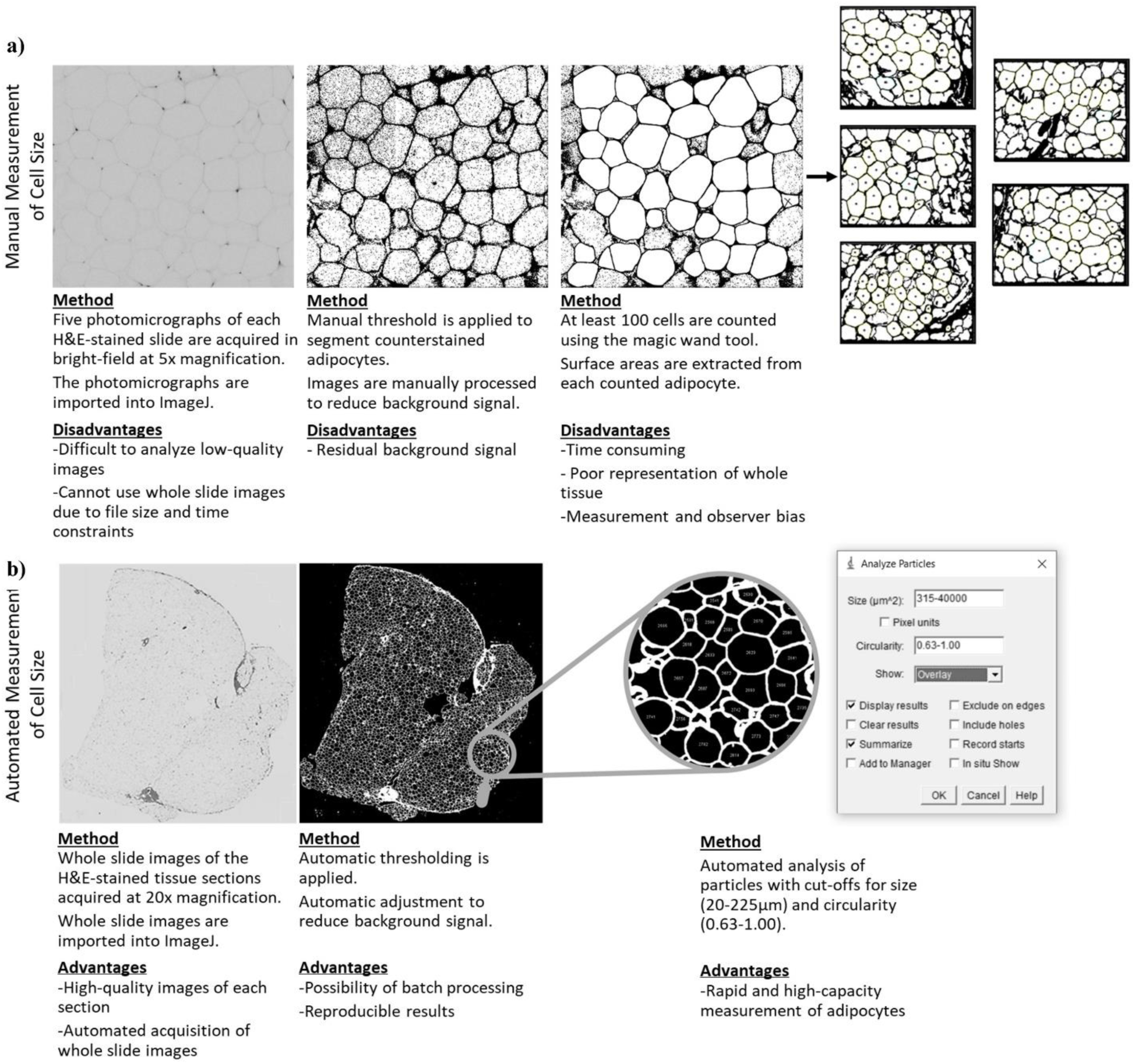
Schematic overview of the manual **(a)** and the automated **(b)** adipocyte diameter measurements in image J. Descriptive steps with disadvantages and advantages of each method are provided.

### 4.4 Statistical analyses

Spearman correlations and paired t-tests were used for the initial validation with 16 adipose tissue samples with GraphPad Prism 10.0.3 (GraphPad Software, LLC, San Diego, Calif, USA).

For the final validation of the automated method using 377 adipose tissue samples from 190 patients, the strength of association between the mean adipocyte diameters from each method was analyzed using paired t-tests and Pearson correlations. Disagreement between the mean adipocyte diameters from each method was evaluated using Bland-Altman plots. Pearson correlations were used to analyze the association between mean adipocyte diameters from each method with different markers of cardiometabolic health (BMI, WC, total Chol, HDL-Chol, LDL-Chol, TG, apoB, HOMA-IR, and VAI). Meng’s Z-test for correlated correlation coefficients was used to determine if the correlations differed significantly based on the method used. Statistical analysis was performed using RStudio® 2024.04.2 Build 764 (Posit Software, PBC, Boston, Mass. USA) with R version 4.3.3 and with GraphPad Prism® Version 10.3.1 (509) (GraphPad Software, LLC, San Diego, Calif, USA).

## Authors’ contribution

Conceptualization: MFG, AR, and AT. Methodology: MFG and AR. Software: MFG. Validation: AR, IMP, MFG, and AT. Formal analysis and Investigation: AR, IMP, and MFG. Resources: AT, FH, LB, and MFG. Data Curation: IMP, GO, AR, and MFG. Supervision: AT. Writing - Original Draft: AR, MFG, and AT. Writing - Review & Editing: AR, MFG, IMP, GO, FH, LB, and AT. Visualization: AR and IMP. Supervision: AT. Project Administration: AT, MFG, and AT Funding acquisition: AT. All authors have read and approved of the final work.

## Data availability statement

The participants of this study did not give written consent for their data to be shared publicly, so due to the sensitive nature of the research supporting data is not available.

